# Subject-specific features of excitation/inhibition profiles in neurodegenerative diseases

**DOI:** 10.1101/2021.12.23.473997

**Authors:** Anita Monteverdi, Fulvia Palesi, Alfredo Costa, Paolo Vitali, Anna Pichiecchio, Matteo Cotta Ramusino, Sara Bernini, Viktor Jirsa, Claudia A.M. Gandini Wheeler-Kingshott, Egidio D’Angelo

## Abstract

Brain pathologies are based on microscopic changes in neurons and synapses that reverberate into large scale networks altering brain dynamics and functional states. An important yet unresolved issue concerns the impact of patients excitation/inhibition profiles on neurodegenerative diseases including Alzheimer’s disease, Frontotemporal Dementia and Amyotrophic Lateral Sclerosis. In this work we used a simulation platform, The Virtual Brain, to simulate brain dynamics in healthy controls and in Alzheimer’s disease, Frontotemporal Dementia and Amyotrophic Lateral Sclerosis patients. The brain connectome and functional connectivity were extracted from 3T-MRI scans and The Virtual Brain nodes were represented by a Wong-Wang neural mass model endowing an explicit representation of the excitatory/inhibitory balance. The integration of cerebro-cerebellar loops improved the correlation between experimental and simulated functional connectivity, and hence The Virtual Brain predictive power, in all pathological conditions. The Virtual Brain biophysical parameters differed between clinical phenotypes, predicting higher global coupling and inhibition in Alzheimer’s disease and stronger NMDA (N-methyl-D-aspartate) receptor-dependent excitation in Amyotrophic Lateral Sclerosis. These physio-pathological parameters allowed an advanced analysis of patients’ state. In backward regressions, The Virtual Brain parameters significantly contributed to explain the variation of neuropsychological scores and, in discriminant analysis, the combination of The Virtual Brain parameters and neuropsychological scores significantly improved discriminative power between clinical conditions. Eventually, cluster analysis provided a unique description of the excitatory/inhibitory balance in individual patients. In aggregate, The Virtual Brain simulations reveal differences in the excitatory/inhibitory balance of individual patients that, combined with cognitive assessment, can promote the personalized diagnosis and therapy of neurodegenerative diseases.

## Introduction

Neuroscience is showing a growing interest in merging results at different scales of complexity in order to achieve a global and comprehensive knowledge of the brain and its mechanisms. In this context, brain modeling can be used to bridge the gap between cellular phenomena and whole-brain dynamics, both in physiological (i.e. healthy) and pathological conditions^1^. The Virtual Brain (TVB)^2,3^ is a neuroinformatic platform recently developed to simulate brain dynamics starting from individual structural (SC) and functional connectivity (FC) matrices constructed from MRI data. TVB has been used to characterize brain dynamics in healthy subjects^4^ but also to explore pathological mechanisms in neurological diseases, such as epilepsy^5^, stroke^6^, brain tumor^7^ and Alzheimer’s disease^8,9^.

Neurodegenerative states ranging from Alzheimer’s disease, Frontotemporal Dementia and Amyotrophic lateral sclerosis are reportedly characterized by a disrupted balance between excitation and inhibition, but this knowledge is not yet available for single subjects unless using lengthy magnetic resonance spectroscopy sequences to quantify Glutamate or GABA (gamma-Aminobutyric acid) concentrations. Hyperexcitation is thought to play a pivotal role in their pathogenesis^10–12^, but multiform and sometimes contradictory results based on empirical observations make it difficult to gain an overall agreement on the neural mechanisms and the evolution of hyperexcitation over the course of the disease. In addition, despite some controversies, increasing findings are supporting GABAergic remodeling as an important feature of Alzheimer’s disease condition^13^. GABAergic dysfunction is less explored in Frontotemporal Dementia and Amyotrophic lateral sclerosis, but it has been demonstrated that baseline GABA levels can influence response to therapies in Frontotemporal Dementia patients^14^ while impaired cortical inhibition due to GABAergic dysfunction can change over Amyotrophic lateral sclerosis progression^15^. Predicting treatment effectiveness for Alzheimer’s disease, Frontotemporal Dementia and Amyotrophic lateral sclerosis patients remains problematic, and the lack of meaningful biomarkers for patients’ classification worsen the situation. Since TVB is designed to extract information about connectivity and network parameters including those linked to inhibition/excitation pathways in single human subjects, it has a high potential to foster personalized and precision medicine.

It is important to point out the need of performing TVB analysis including not only cerebral nodes and their structural connections to one another but also the cerebellum. Recently, it was shown that integrating cerebro-cerebellar connections can improve TVB predictive capability in healthy subjects^16^. This is in line with the increasing evidence supporting cerebellar involvement not only in motor learning and coordination^17^ but also in cognitive processing^18,19,20,21^. Cerebellar impairment has been revealed in Alzheimer’s disease^22–24^, Frontotemporal Dementia^25^ and Amyotrophic lateral sclerosis^26^.

In this work, we exploited TVB capabilities i) to characterize each group of subjects by providing personalized excitation/inhibition profiles ii) to assess the cerebellar impact on brain dynamics generation in healthy controls and in Alzheimer’s disease, Frontotemporal Dementia and Amyotrophic lateral sclerosis. TVB simulations were performed using the Wong-Wang model^27^, which allowed us to derive a set of subject-specific biophysical parameters able to describe global brain dynamics and the excitatory/inhibitory balance in local networks. We evaluated the potential for clinical translation of the biophysical parameters obtained from TVB simulations by exploring their association with patients’ cognitive performance and testing their discriminative power between clinical conditions and neuropsychological domains. This work, by providing a unique description of the excitatory/inhibitory balance at single-subject level, can contribute to the progress of personalized and precision medicine opening new perspectives for brain modelling in neurodegenerative diseases.

## Materials and Methods

In this work individual’s subject analysis was conducted as described in Figure 1 and simulations were performed in three networks^16^: whole-brain network, cortical subnetwork, and embedded cerebro-cerebellar subnetwork (see section networks, Figure 2).

**Figure 1.**
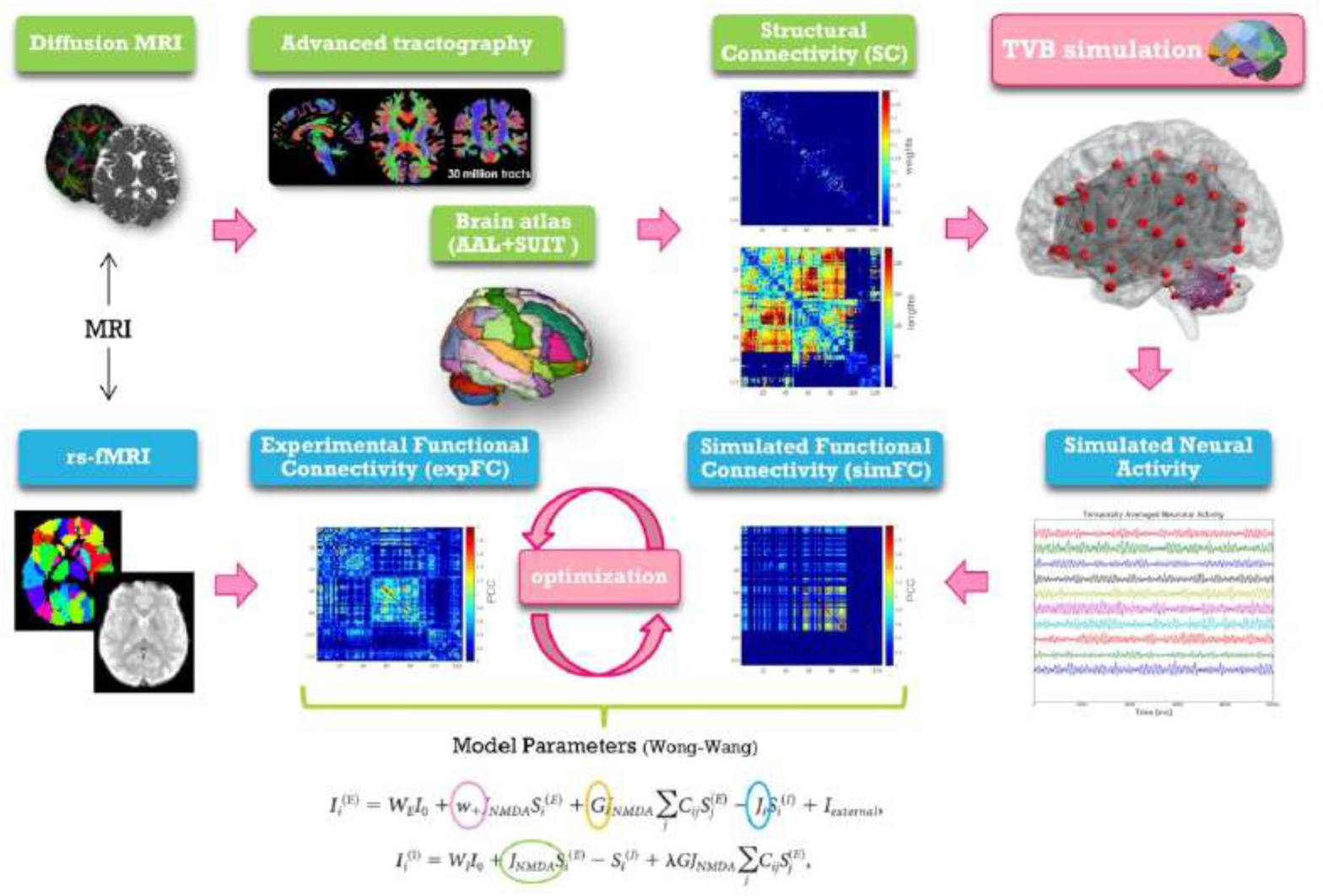
Schematic representation of modelling workflow. MRI is used to obtain *the structural and experimental functional connectivity matrices* needed for TVB construction and optimization. From top left, clockwise: diffusion weighted images are preprocessed and elaborated to yield wholebrain tractography. An ad-hoc parcellation atlas combining AAL atlas and SUIT is used to map the structural connectivity (SC) matrices obtained from whole-brain tractography parcellation (top weight matrix, bottom distance matrix). The Virtual Brain (TVB) is constructed using the structural connectivity matrix for edges and neural masses for nodes. TVB simulations of neural activity allow to extract BOLD signals for each node leading to define the *simulated functional connectivity* (FC) *matrix*. TVB optimization is performed through model inversion by comparing the simulated FC with the experimental FC. Model parameters and the corresponding equations are shown at the bottom.

**Figure 2.**
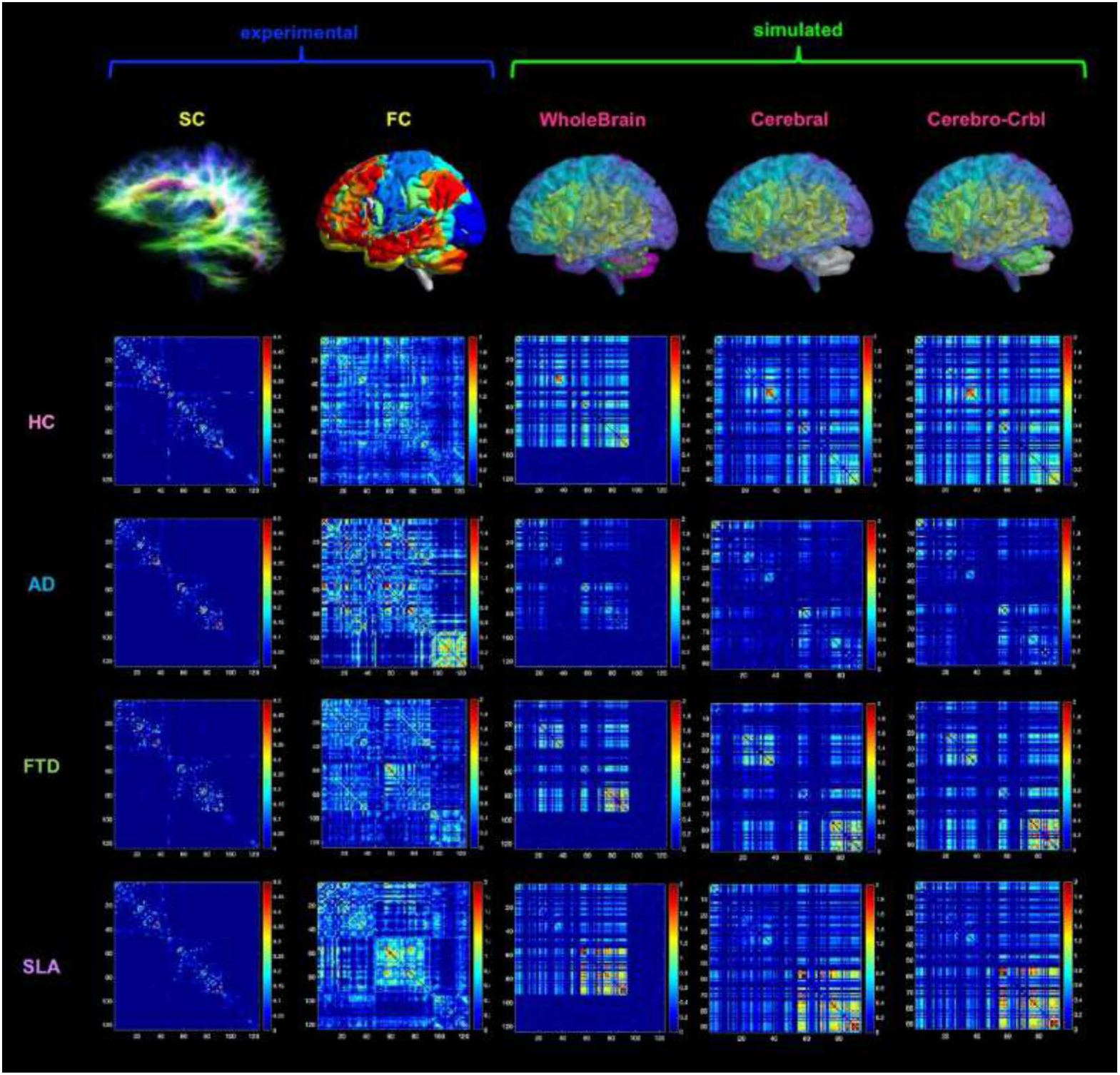
Networks connectivity matrices. Columns 1 and 2 show the experimental structural (SC) and functional connectivity (FC) matrices, which were used as input for TVB simulations in four different groups: healthy (HC), Alzheimer’s disease (AD), Frontotemporal Dementia (FTD) and Amyotrophic Lateral Sclerosis (ALS). For each group, matrices of a randomly chosen subject are reported as an example. Columns 3 to 5 show the simulated FC obtained at single-subject level with three different networks: whole-brain, cortical subnetwork (Cerebral) and embedded cerebro-cerebellar subnetwork (Cerebro-Crbl). In the whole-brain network simulations were performed using whole-brain nodes and connections (whole-brain nodes and edges are colored); in the cortical subnetwork only cerebral cortex nodes and connections were considered (cortical nodes and edges are colored); in the embedded cerebro-cerebellar subnetwork cerebral cortex nodes were considered taking into account the influence of cerebro-cerebellar connections (cortical nodes and cerebellar edges are colored).

### Subjects

Sixty patients affected by neurodegenerative diseases were recruited at the IRCCS Mondino Foundation, as part of a study on cognitive impairment published in Palesi et al.^24^, Castellazzi et al.^28^, Lorenzi et al.^29^, Pizzarotti et al.^25^. The study was carried out in accordance with the Declaration of Helsinki with written informed consent from all subjects. The protocol was approved by the local ethic committee of the IRCCS Mondino Foundation. Patients underwent a complete diagnostic workup including neuropsychological assessment, MRI (and electroneuromyography in patients with motor neuron impairment) in order to obtain an exhaustive phenotypic profiling and a correct etiological definition. Based on the most recent diagnostic criteria subjects were classified into three groups: fifteen Alzheimer’s disease patients^30^ (6 females, 70 ± 7 years), fifteen Frontotemporal Dementia patients (4 females, 69 ± 7 years) [including behavioral Frontotemporal Dementia ^31^ and Primary Progressive Aphasia^32^], fifteen Amyotrophic lateral sclerosis^33^ patients (7 females, 67 ± 8 years). In detail, diagnosis of Alzheimer’s disease was made according to the criteria of the National Institute of Neurological and Communicative Disorders and Stroke and Alzheimer’s Disease and Related Disorders Association (NINCDS-ADRDA) workgroup^30^; Frontotemporal Dementia diagnosis was supported according to Rascovsky diagnostic criteria but not determined by the cognitive profile and no patient was excluded based on neuropsychological profile if diagnostic criteria were still met; Amyotrophic lateral sclerosis diagnosis was made in patients fulfilling Awaji criteria^33^ and this group included patients with Amyotrophic lateral sclerosis and mild cognitive impairment. In addition, fifteen healthy controls (8 females, 64 ± 11 years) were enrolled on a voluntary basis as reference group. All healthy controls underwent clinical assessment to exclude any cognitive or motoneuron impairment.

For all subjects, exclusion criteria were: age>80 years, a diagnosis of significant medical, neurological (other than Alzheimer’s disease, Frontotemporal Dementia, Amyotrophic lateral sclerosis) and psychiatric disorder, pharmacologically treated delirium or hallucinations and secondary causes of cognitive decline (e.g. vascular metabolic, endocrine, toxic and iatrogenic). Table 1 shows demographic, clinical, and neuropsychological data.

**Table 1:** Demographics, clinical and neuropsychological data (means, SDs and group differences).

| Measures | HC | AD | FTD | ALS | p-value |
| --- | --- | --- | --- | --- | --- |
| Males/females | 7/8 | 9/6 | 11/4 | 8/7 | 0.201 |
| Age (years) | 64 (11) | 70 (7) | 69 (7) | 67 (8) | 0.494 |
| Education (years) | 10 (3) | 8 (3) | 10 (4) | 9 (5) | - |
| Memory | 3.0 (0.4) | 0.7 (0.7) | 1.6 (0.7) | 2.6 (0.6) | <0.001 * |
| Executive-function | 2.9 (0.6) | 0.8 (1.1) | 1.0 (0.9) | 1.9 (0.9) | <0.001 * |
| Attention | 3.4 (0.5) | 1.1 (1.0) | 1.3 (1.2) | 2.0 (0.8) | <0.001 * |
| Language | 3.5 (0.5) | 1.1 (1.3) | 1.3 (1.0) | 2.9 (1.2) | <0.001 * |
| Visuospatial skills | 3.7 (0.8) | 1.2 (1.7) | 2.2 (1.9) | 2.2 (2.0) | 0.002 * |

Gender, age, education and neuropsychological scores are reported for each group (Healthy controls, HC, Alzheimer's disease, AD, Frontotemporal dementia, FTD and Amyotrophic lateral sclerosis, ALS) as mean values and standard deviations in brackets. Significant threshold is set at $p < 0.05$ .
\* refers to significant group differences assessed with Kruskal-Wallis

### Neuropsychological examination

All subjects underwent a neuropsychological examination based on a standardized battery of tests to assess their global cognitive status (Mini-Mental State Examination, MMSE^34^) and different cognitive domains: attention (Stroop test^35^, Trail Making test ^36^A and B, Attentive Matrices^37^), memory (Digit and Verbal span, Corsi block-tapping test, Logical Memory test^37^, Rey-Osterrieth^38^ complex figure delayed recall, Rey’s 15 words test^39^), language (phonological^39^ and semantic^40^verbal fluency), logical-executive functions (Raven’s Matrices^39^ 1947, Winconsing Card Sorting^41^ test, Frontal Assessment^42^ Battery) and visuospatial skills (Rey-Osterrieth^38^ complex figure copy). For each test age-, gender- and education corrected-scores were computed and then transformed into equivalent scores ranging from 0 (pathological) to 4 (normal) on the basis of the equivalent score standardization method^43^. For each cognitive domain, a weighted score was derived from the average of the equivalent scores of the tests belonging to that specific cognitive domain.

### MRI Acquisitions

All subjects underwent MRI examination using a 3T Siemens Skyra scanner with a 32-channel head coil. The protocol included resting-state fMRI (T_2_*-weighted GRE-EPI sequence, TR/TE = 3010/20 ms; 60 slices, acquisition matrix = 90×90, voxel size = 2.5×2.5×2.5 mm^3^ isotropic, 120 volumes) and diffusion weighted (DW) imaging (SE-EPI sequence, TR/TE = 10000/97 ms, 70 slices with no gap, acquisition matrix = 120×120, voxel size = 2×2×2 mm^3^ isotropic, 64 diffusion-weighted directions, b-value = 1200 s/mm^2^, 10 volumes with no diffusion weighting (bo image). For anatomical reference, a whole brain high-resolution 3D sagittal T1-weighted (3DT1) scan (TR/TE = 2300/2.95 ms, TI = 900 ms, flip angle = 9°, 176 slices, acquisition matrix = 256 × 256, in-plane resolution = 1.05 × 1.05 mm, slice thickness = 1.2 mm) was also acquired.

### Preprocessing and tractography of diffusion data

For each subject, a mean b_0_ image was obtained averaging the 10 volumes acquired with no diffusion weighting. DW data were denoised and corrected for Gibbs artifact^44^, eddy currents distortions and aligned to the mean bo image using eddy tool^45^ (FSL). A binary brain mask was obtained from the mean b_0_ image using brain extraction tool^46^ and DTIFIT was used to generate individual fractional anisotropy (FA) and mean diffusivity (MD) maps. 3DT1-weighted images were segmented using MRtrix3^47,48^ as white matter (WM), gray matter (GM), subcortical GM and cerebrospinal fluid (CSF). 30 million streamlines whole-brain anatomically constrained tractography^49^ was performed within MRtrix3, estimating fibers orientation distribution with multi-shell multi-tissue constrained spherical deconvolution (CSD) and using probabilistic streamline tractography^50^. As in previous works^16,51^, a correction of spurious cerebro-cerebellar tracts was performed excluding the ipsilateral connections from whole-brain tractograms.

### Preprocessing of fMRI data

fMRI preprocessing was carried out combining SPM12^52^, FSL and MRtrix3 commands in a custom MATLABR2019b^53^ script. Marchenko-Pastur principal component analysis (MP-PCA) denoising^54^ was firstly performed, followed by slice-timing correction, realignment to mean functional image and affine registration to the 3DT1-weighted image. These steps were followed by polynomial detrend and 24 motion parameters regression^55^. A subject-specific CSF mask was extracted from the 3DT1 segmentation, eroded using a 99% probability threshold, and constrained to areas within the ALVIN (Automatic Lateral Ventricle delIneatioN) mask of the ventricles^56^. These corrections were performed to avoid the risk of capturing signals of interest from adjacent GM voxels, and nuisance regressors identified within the restricted CSF mask were removed using a component-based noise correction (compCor) approach^57,58^. Temporal band-pass filtering (0.008-0.09 Hz) was finally applied.

### Structural and functional connectivity

Connectomes of SC and FC were estimated combining a parcellation atlas with whole-brain tractography and rs-fMRI signals of each subject, respectively. An ad-hoc GM parcellation atlas was created combining 93 cerebral (including cortical and deep GM structures) and 31 cerebellar (SUIT, A spatially unbiased atlas template of the cerebellum and brainstem) labels^59^ in MNI152 space. Each GM parcellation was considered as a node for the connectivity analysis. The atlas was transformed to subject-space inverting the normalization from the 3DT1-weighted image to the MNI152 standard space. The parcellation atlas applied to the whole-brain tractography led to two types of SC matrices: a distance matrix containing the length of tracts connecting each pair of nodes, and a weight matrix in which connections strengths (number of streamlines) were normalized by the maximum value per each subject. The time-course of BOLD signals was extracted for each node and the experimental FC matrix (expFC) was computed as the Pearson’s correlation coefficient (PCC) of the time-courses between each pair of brain regions. Matrix elements were converted with a Fisher’s z transformation and thresholded at 0.1206^16^.

### Brain dynamics simulation with TVB

TVB workflow includes several steps: 1) incorporation of subject SC matrices; 2) selection of a mean field/neural mass mathematical model; 3) simulation of the rs-fMRI time-courses per node and creation of the simulated FC matrix (simFC); 4) model parameters tuning to achieve the best matching between simFC and expFC matrices; 5) final simulation of brain dynamics with the optimal model parameters as described in detail by Deco *et al*.^27^.

### Computational model from neuronal activity to large-scale signals

The Wong-Wang model^27^ implemented as highly optimized C code^60^ was chosen to simulate wholebrain dynamics. This dynamic mean field model simulates the local regional neuronal activity as the result of a network composed of interconnected excitatory and inhibitory neurons coupled through NMDA and GABA receptor types. Details on the Wong-Wang model can be found in Deco *et al*.^27^. Briefly, brain dynamics are described by the following set of coupled non-linear stochastic differential equations:

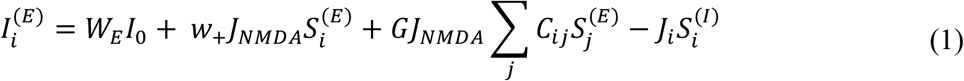

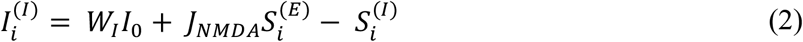

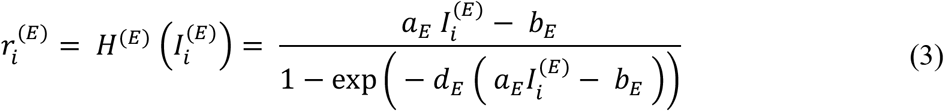

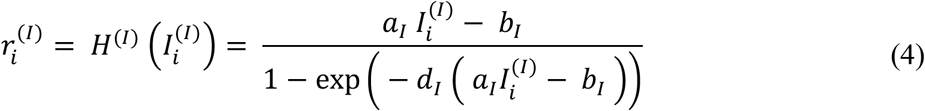

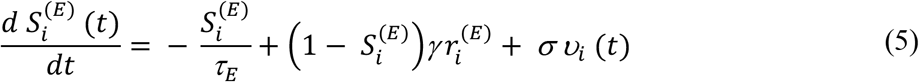

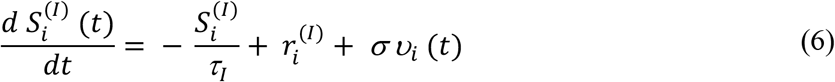

where r_i_^(E,I)^ denotes the firing rate of the excitatory (E) and inhibitory (I) population, S_i_^(E,I)^ identifies the average excitatory or inhibitory synaptic gating variables at each local area, i, and I_i_^(E,I)^ is the input current to the excitatory and inhibitory populations. All parameters described in Supplementary Table 1 were set as in Deco *et al*.^27^, except those that are tuned during parameters optimization, described below. Parameter space exploration was performed for global coupling (G), which is a scaling factor denoting long-range coupling strength, and local parameters defining the strength of inhibitory (GABA) synapses (J_i_), the strength of excitatory (NMDA) synapses (J_NMDA_) and the strength of local excitatory recurrence (w_+_). Thus, this model retains information both on global brain dynamics and local excitatory/inhibitory balance and is particularly interesting for the investigation of pathological conditions.

For each set of parameters combination, resting-state BOLD fMRI time-courses were simulated over 6 min length using a Balloon-Windkessel hemodynamic neurovascular coupling model^61^ while the simFC was computed as described below (see section 2.7) for the expFC. Parameters were adjusted iteratively until the best fit, i.e. the highest correlation, between expFC and simFC was achieved (Supplementary Figure 1).

### Networks

To investigate the impact of the cerebellum on brain dynamics generation, simulations were performed using three different combinations of connections and nodes (Figure 2):

- Whole-brain network: whole-brain nodes and connections
- Cortical subnetwork: cerebral cortex nodes and connections (excluding cerebro-cerebellar connections)
- Embedded cerebro-cerebellar subnetwork: cerebral cortex nodes but also considering the influence of cerebro-cerebellar connections

For each of these three networks predictive power was evaluated as the mean PCC between expFC and simFC matrices in different clinical conditions (healthy controls, Alzheimer’s disease, Frontotemporal Dementia, Amyotrophic lateral sclerosis).

### Statistic

Statistical tests were performed using SPSS software version 21^62^.

### Excitation/inhibition role in neurodegeneration

To assess whether biophysical parameters derived from TVB differ according to the clinical condition, optimal model parameters were tested for normality (Shapiro-Wilk test) and differences between groups were assessed with non-parametric tests (Kruskal-Wallis across all groups and Mann-Whitney between each pair of groups) because they did not present a Gaussian distribution.

A multiple regression analysis was performed to investigate the relationship between individual scores of the 6 cognitive domains (attention, memory, language, logical-executive functions, visuospatial skills) and the optimal model parameters. Model parameters combined with age, gender and group category were used in a backward approach to identify which of them significantly (p<0.05) explained neuropsychological scores variance in all subjects together.

Moreover, to assess the relevance of these parameters in discriminating between physiological and pathological conditions, a discriminant analysis was performed using the group as the dependent variable and considering as independent variables: (i) model parameters alone, (ii) neuropsychological scores alone and (iii) a combination of both. To visualize and assess the sensitivity and specificity of the best discriminative variables, receiving operating characteristics (ROC) curves and corresponding areas under the curve (AUC) were calculated.

Finally, a k-mean cluster analysis was performed to reconstruct subjects-specific excitation/inhibition profiles. The number of clusters was an input parameter arbitrary set equal to 4 as the number of variables considered (model parameters). A frequency analysis of the clusters obtained in each group enabled a deeper understanding of excitatory/inhibitory balance disruption.

### Cerebellar role in brain dynamics in neurodegeneration

PCC obtained with the three networks were normally distributed (Shapiro-Wilk test), thus parametric tests were used to compare them between different conditions. First, to assess changes of TVB predictive power with the clinical condition, a one-way ANOVA was performed between PCC of each network across groups (healthy controls, Alzheimer’s disease, Frontotemporal Dementia, Amyotrophic lateral sclerosis). Then, to assess the impact of the specific network on TVB predictive power, a multivariate general linear model (GLM) with Bonferroni correction was chosen to compare PCC values of the three networks within each group.

### Code and data accessibility

All codes used for this study are freely available. The optimized TVB C code can be found at https://github.com/BrainModes/fast_tvb. Dataset will be available at 10.5281/zenodo.5796063.

## Results

### Excitation/inhibition role in neurodegeneration

Both global (G) and local (J_i_, J_NMDA_, w_+_) parameters were adjusted iteratively to optimize the model fit to empirical data. Optimal model parameters were found across the whole-brain network of each subject.

### Group differences in TVB parameters

The biophysical parameters derived from TVB were compared between groups to assess whether, at group level, their value could differ according to the clinical condition. Both global and local biophysical parameters showed significant differences between groups (Table 2, Figure 3A): Alzheimer’s disease patients showed higher G and Ji compared to healthy controls and Frontotemporal Dementia (p<0.05); Amyotrophic lateral sclerosis patients showed higher J_NMDA_ than healthy controls (p<0.05); no statistically significant differences were found in healthy controls and Frontotemporal Dementia compared to the other groups.

**Table 2:**
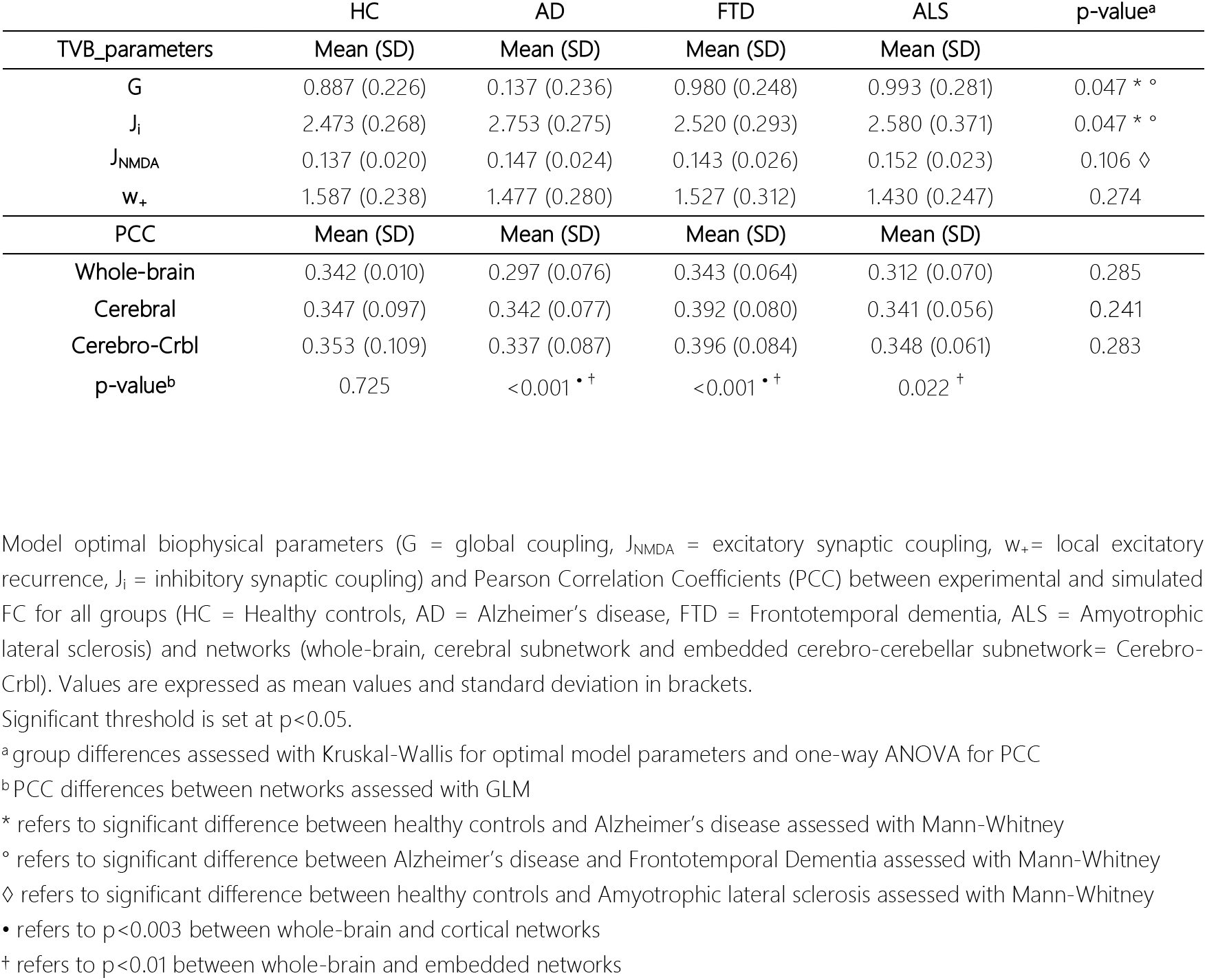
Optimal model parameters and Pearson Correlation Coefficients per group.

**Figure 3.**
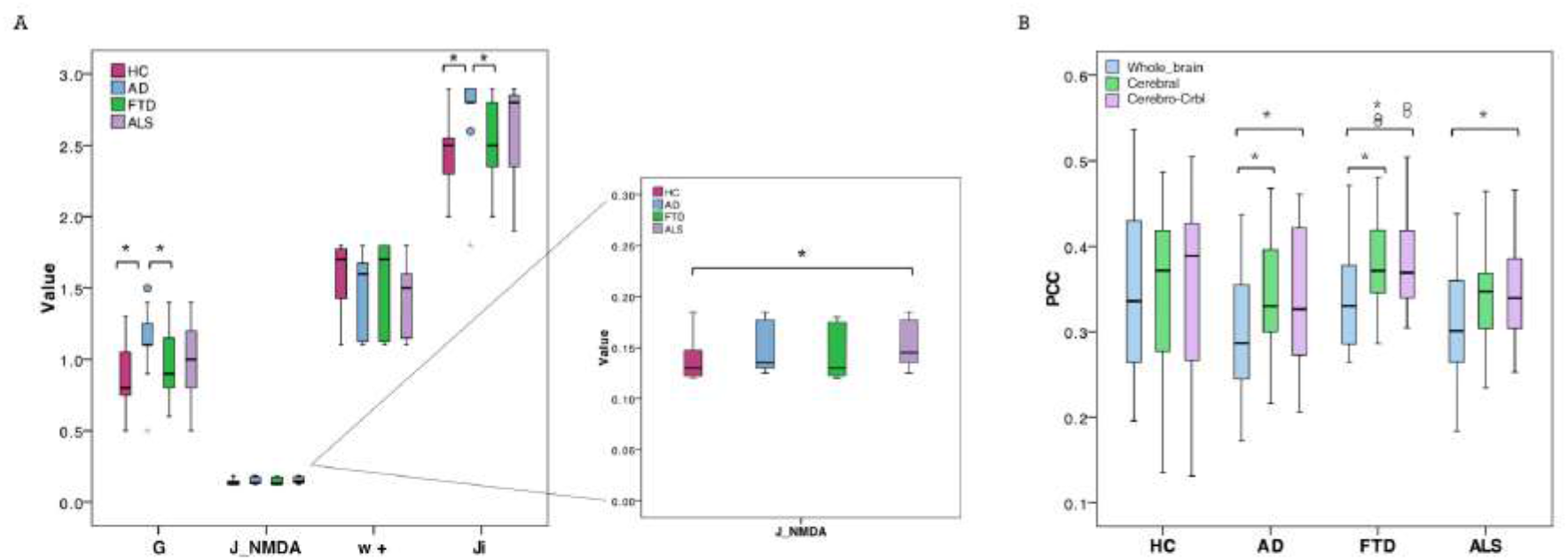
Boxplots of optimal biophysical parameters and Pearson correlation coefficients (PCC). **(A)** Boxplots of optimal biophysical parameters derived from TVB (global coupling, G, excitatory synaptic coupling, J_NMDA, local excitatory recurrence, wp, inhibitory synaptic coupling, Ji) across groups (healthy controls, Alzheimer’s disease, Frontotemporal Dementia and Amyotrophic lateral sclerosis). The asterisk (*) indicates significant difference (Mann-Whitney, p<0.05) between groups (see Table 2 for details). **(B)** Boxplots of Pearson correlation coefficients (PCC) between experimental and simulated FC for all groups (healthy controls, Alzheimer’s disease, Frontotemporal Dementia, Amyotrophic lateral sclerosis) and networks (whole-brain network, Whole_brain, cortical subnetwork, Cerebral; embedded cerebro-cerebellar subnetwork, Cerebro-Crbl). Asterisks (*) indicate significant difference (p<0.05) between networks (see Table 2 for details).

### Relationship between TVB parameters and neuropsychological scores

Parameters used in backward regressions, significantly explained the variation of scores in different neuropsychological domains. The explained variance of each neuropsychological domain was progressively reduced by simplifying the regression model through the removal of one or more predictors and ranged from ~20% to ~8%. For each cognitive domain, a different combination of features was necessary to significantly (p<0.05) explain a percentage of the variance (Table 3).

**Table 3:**
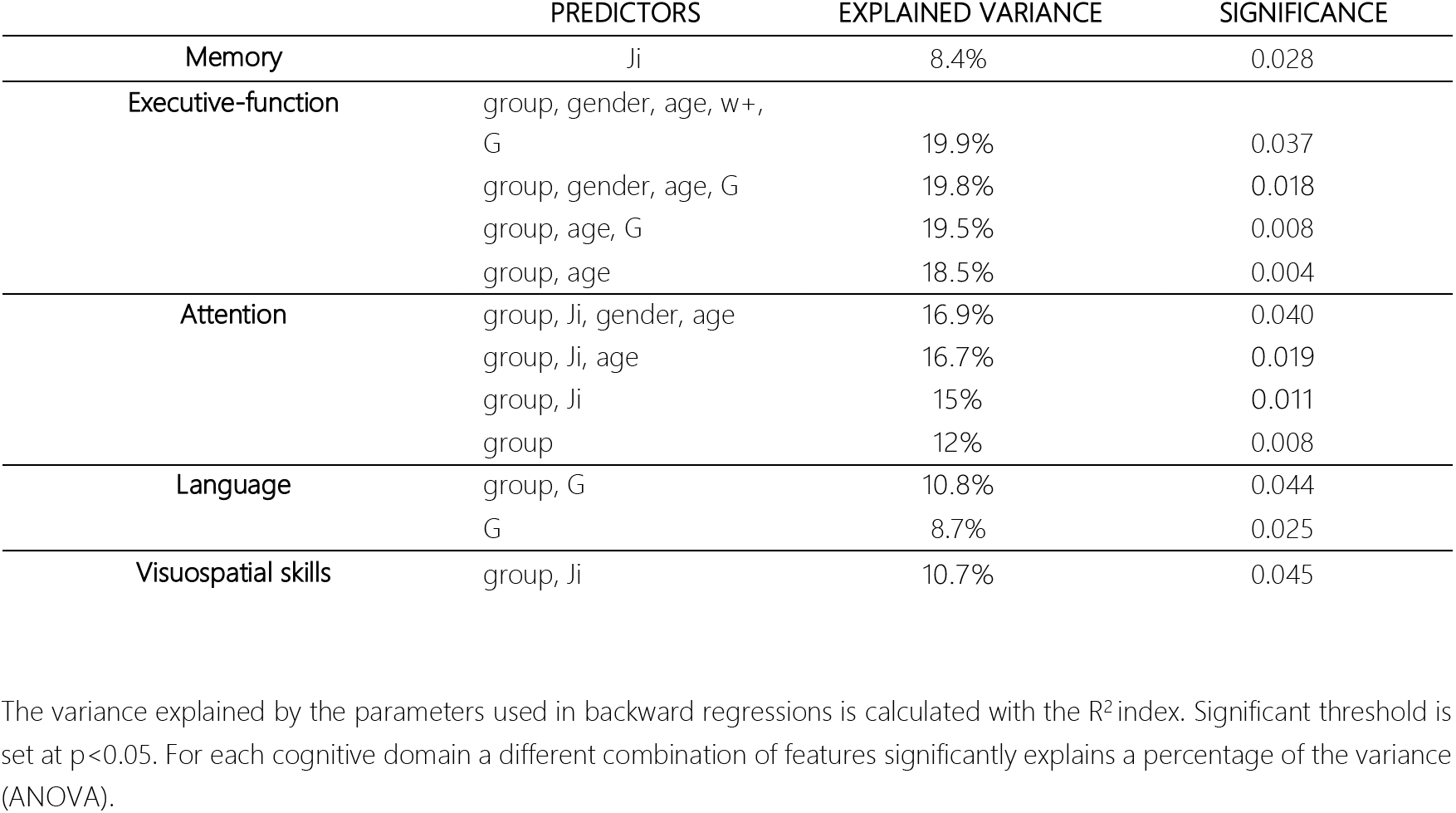
Backward regressions results.

|  | PREDICTORS | EXPLAINED VARIANCE | SIGNIFICANCE |
| --- | --- | --- | --- |
| Memory | Ji | 8.4% | 0.028 |
| Executive-function | group, gender, age, w+, G | 19.9% | 0.037 |
|  | group, gender, age, G | 19.8% | 0.018 |
|  | group, age, G | 19.5% | 0.008 |
|  | group, age | 18.5% | 0.004 |
| Attention | group, Ji, gender, age | 16.9% | 0.040 |
|  | group, Ji, age | 16.7% | 0.019 |
|  | group, Ji | 15% | 0.011 |
|  | group | 12% | 0.008 |
| Language | group, G | 10.8% | 0.044 |
|  | G | 8.7% | 0.025 |
| Visuospatial skills | group, Ji | 10.7% | 0.045 |

The variance explained by the parameters used in backward regressions is calculated with the $R^2$ index. Significant threshold is set at $p < 0.05$ . For each cognitive domain a different combination of features significantly explains a percentage of the variance (ANOVA).

### Discriminative power of TVB parameters and neuropsychological scores

Discriminative power of TVB parameters and neuropsychological scores is reported for all comparisons (healthy controls versus Alzheimer’s disease, healthy controls versus Frontotemporal Dementia, healthy controls versus Amyotrophic lateral sclerosis, Alzheimer’s disease versus Frontotemporal Dementia, Alzheimer’s disease versus Amyotrophic lateral sclerosis, Frontotemporal Dementia versus Amyotrophic lateral sclerosis) in Supplementary Table 2. TVB parameters always yielded a poorer discriminant power (about 70%) than that offered by neuropsychological scores (about 90%). When neuropsychological scores were complemented by TVB values as joint independent variables, the discriminative power increased in all classifications reaching 100% when distinguishing between Alzheimer’s disease and healthy controls, and between Frontotemporal Dementia and Amyotrophic lateral sclerosis. To visualize all these results, ROC curves are reported in Figure 4.

**Figure 4.**
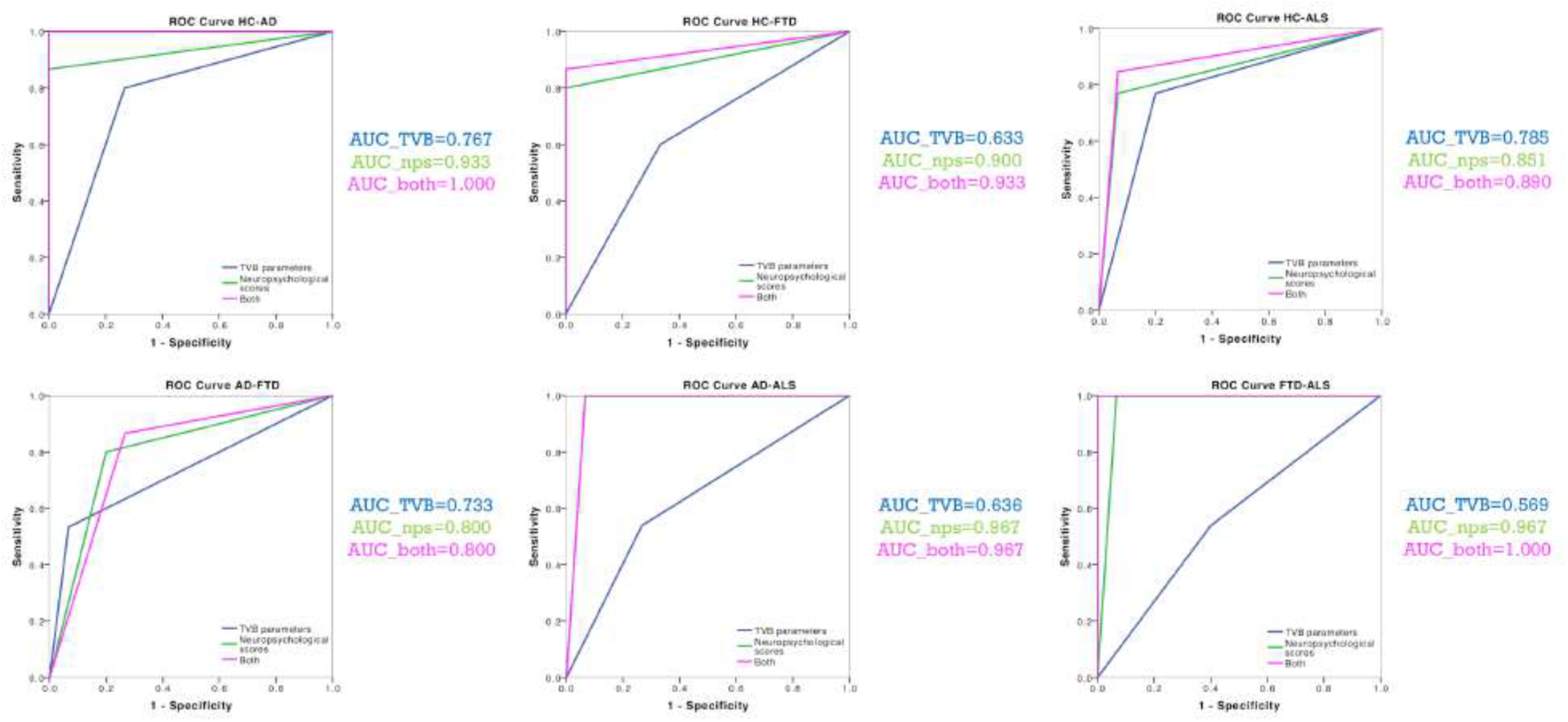
Classification analysis. ROC curves were calculated for each classification (healthy controls versus Alzheimer’s disease, healthy controls versus Frontotemporal Dementia, healthy controls versus Amyotrophic lateral sclerosis, Alzheimer’s disease versus Frontotemporal Dementia, Alzheimer’s disease versus Amyotrophic lateral sclerosis, Frontotemporal Dementia versus Amyotrophic lateral sclerosis) with their corresponding groups of variables (TVB parameters alone, neuropsychological scores alone, TVB parameters combined with neuropsychological scores). AUC values confirm that TVB parameters alone (blu) always yielded a poorer discriminant power than that offered by neuropsychological scores alone (green). The combination of TVB parameters with neuropsychological scores improved the discriminative power in all classifications reaching 100% when distinguishing between Alzheimer’s disease and healthy controls and between Frontotemporal Dementia and Amyotrophic lateral sclerosis.

### A personalized description of the excitatory/inhibitory balance

Each of the four clusters identified with the k-means analysis was characterized by a different combination of values for TVB-derived biophysical parameters, as reported in Figure 5A and Supplementary Figure 2. Considering the biophysical meaning of each parameter derived from the simulation, we can describe the k-means clusters as follows:

- cluster 1 is mainly characterized by medium to strong overexcitation (medium to high values of J_NMDA_)
- cluster 2, in addition to show strong overexcitation (very high values of J_NMDA_), is characterized by a high global coupling strength (medium to high values of G) and medium to strong overinhibition (medium to high values of J_i_)
- cluster 3 is the only one characterized by medium or low values of G, J_i_ and J_NMDA_ and high values of local excitatory recurrence (w_+_)
- cluster 4 is mostly characterized by overinhibition (high values of J_i_) and high global coupling strength between nodes (higher values of G).

**Figure 5.**
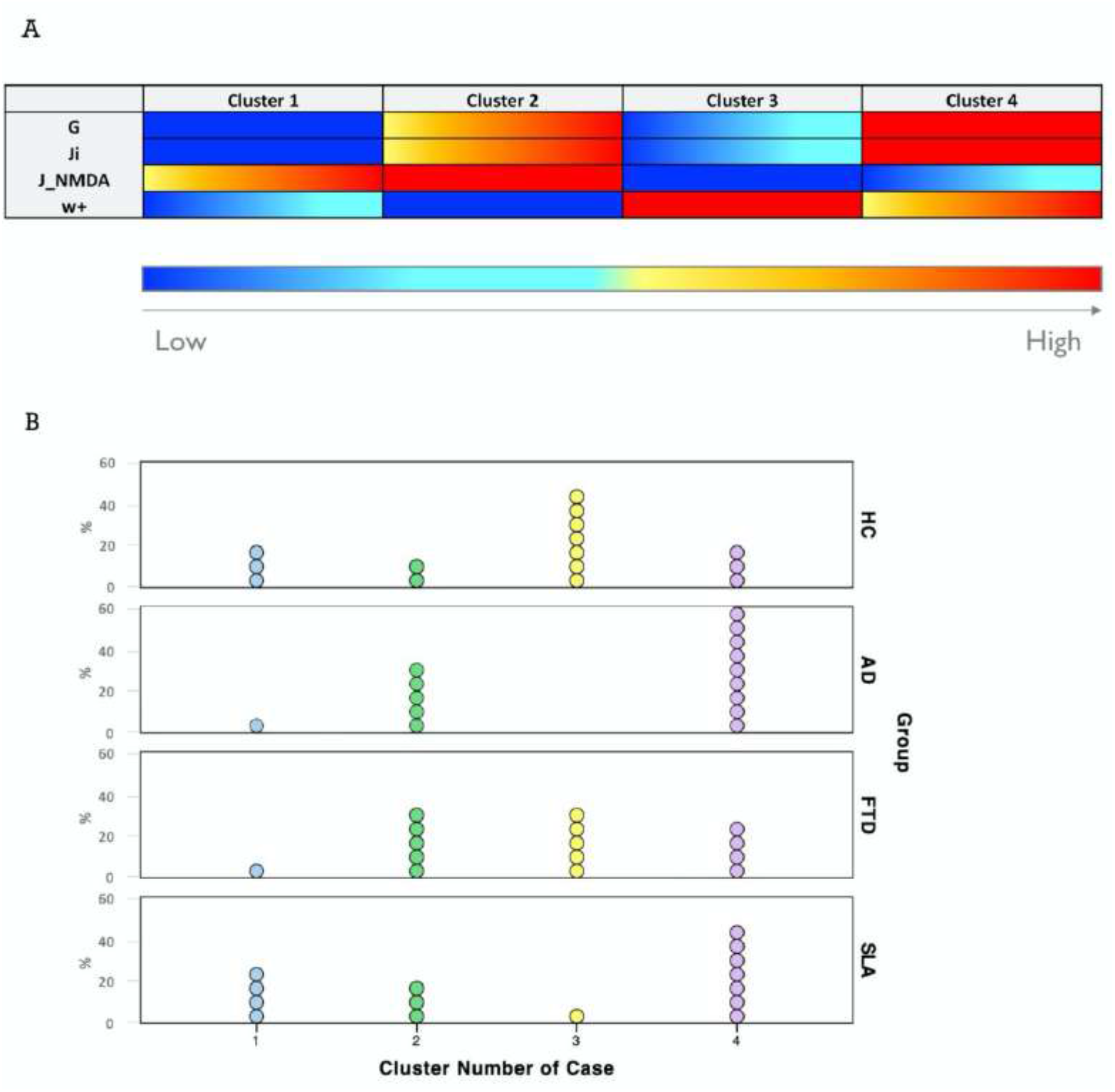
Excitation/inhibition profiles. **(A)** Each cluster was characterized by a typical excitation/inhibition profile. The color-bar (from blue to red) represents the scale from low to high of each TVB-derived biophysical parameter. (**B)** Visual representation of cluster distributions across groups (healthy controls, Alzheimer’s disease, Frontotemporal Dementia and Amyotrophic lateral sclerosis). Cluster numbers are reported on the x axis while cluster frequencies in each condition are reported on the y axis. Each dot represents a single subject.

All the groups considered were represented in more than one clusters, as shown in Figure 5B. The distribution of subjects belonging to different condition across clusters revealed that, cluster 1 was more frequent in healthy controls (20%) and Amyotrophic lateral sclerosis (26.7%) than in Alzheimer’s disease and Frontotemporal Dementia (both 6.7%), cluster 2 was less present in healthy controls (13.3%) than in pathological conditions (Alzheimer’s disease=33.3%, Frontotemporal Dementia=33.3% and Amyotrophic lateral sclerosis=20%), cluster 3 was the mostly present in healthy controls subjects (46.7%), medium frequent in Frontotemporal Dementia (33.3%) and tended to disappear in Amyotrophic lateral sclerosis (6.7%) and Alzheimer’s disease subjects (0%), cluster 4 was the most frequent in Alzheimer’s disease (60%) and Amyotrophic lateral sclerosis (46.7%) and less frequent in healthy controls (20%) and Frontotemporal Dementia (26.7%).

### Cerebellar role in brain dynamics in neurodegeneration

To understand the role of the cerebellum in neurodegeneration, we performed TVB simulations using three different networks: (i) Whole-brain network, (ii) Cortical subnetwork, (iii) Embedded cerebro-cerebellar subnetwork as in Palesi *et al*.^16^. For each of these three networks, the predictive power was evaluated as the mean PCC between expFC and simFC matrices in healthy controls and in the pathological groups: Alzheimer’s disease, Frontotemporal Dementia, Amyotrophic lateral sclerosis.

### Predictive power of TVB simulations

TVB simulation performed both in physiological and pathological conditions led to good fit values between the expFC and simFC (Table 2). No differences were found between PCC values of each network across clinical groups, but significant differences were found in each group comparing PCC of the three networks. For all the pathological groups, PCC values obtained with the embedded cerebro-cerebellar subnetwork were significantly higher (p<0.01) than those obtained with the wholebrain network, while PCC of the cortical subnetwork was significantly higher (p<0.003) than that of the whole-brain network in Alzheimer’s disease and Frontotemporal Dementia (Table 2, Figure 3B). No differences were found between the three networks in healthy controls.

## Discussion

For the first time, in this work we characterized the excitatory/inhibitory profile in neurodegenerative (Alzheimer’s disease, Frontotemporal Dementia, Amyotrophic lateral sclerosis) conditions integrating cerebro-cerebellar connections in TVB. Importantly, by adopting the Wong-Wang model to model brain dynamics, we gained information on local excitatory/inhibitory balance at the single subject level.

### Excitation/inhibition role in neurodegeneration

#### Hyper-excitation and over-inhibition underly different neurodegenerative mechanisms

Parameters derived from TVB simulations using the Wong-Wang model yield information on global brain dynamics and local excitatory/inhibitory balance. In particular, the global scaling factor G denotes the strength of long-range connections, and higher global coupling means a greater weighting of the global over the local connectivity. The remaining three parameters define the balance between excitation and inhibition in the simulated network: J_NMDA_ represents the strength of excitatory synapses in the network, J_i_ denotes the strength of inhibitory synapses and w_+_ the strength of recurrent excitation.

Our results revealed that different clinical groups were characterized by specific TVB parameters providing new clues for the interpretation of the dysfunctional mechanisms in local microcircuits.

To date, patterns of altered FC in Alzheimer’s disease from a resting-state networks perspective have been reported mainly in the default mode network (DMN)^63,64^, although a wider involvement has been suggested by our group in previous works^22^. Our data demonstrated that G and J_i_ were higher in Alzheimer’s disease patients compared to healthy controls suggesting that Alzheimer’s disease subjects were characterized by increased global coupling and overinhibition. This increased G value in our Alzheimer’s disease group could be interpreted as a compensatory mechanism counteracting altered cerebral connectivity, but it might also underly the hypersynchrony typically characterizing disrupted networks in patients^22,65^. Furthermore, our data showing an increased inhibition and suggesting that GABAergic dysfunction plays a role in Alzheimer’s disease pathology are in line with the novel hypothesis that GABAergic remodeling might be an important feature of neurodegeneration^13^.

It is worth noting that Alzheimer’s disease showed higher G and J_i_ also compared to Frontotemporal Dementia patients, strengthening the tenet that the pathophysiological mechanisms underlying the two diseases are different. Indeed, Frontotemporal Dementia showed G and J_i_ values similar to healthy controls, consistent with the similarity of cortical neural synchronization in Frontotemporal Dementia and healthy controls^66^.

Finally, Amyotrophic lateral sclerosis patients were characterized by an increased J_NMDA_, which is in line with the cortical hyperexcitability frequently reported in this pathological condition^12^.

#### TVB-derived biophysical parameters help to explain cognitive performance

TVB-derived biophysical parameters combined with age, gender and group category significantly explained the variance of neuropsychological scores both in physiological and pathological groups. This suggests that the levels of excitation, inhibition and global coupling are associated with cognitive performance in the different clinical conditions.

The clinical relevance of TVB parameters is further highlighted by results of discriminant analysis. The discriminative power of neuropsychological tests alone was always higher than the one obtained with TVB parameters alone. However, when neuropsychological measures were combined with TVB parameters, the discriminative power improved, reaching in some cases 100% of accuracy. Importantly the performance of our classification was satisfactory not only to distinguish healthy controls from patients, but also to differentiate patients belonging to different neurodegenerative conditions. This opens an interesting perspective for the development of new diagnostic tools combining TVB parameters with neuropsychological scores in future machine learning approaches.

#### The excitatory/inhibitory balance in single-subject

For each subject, TVB predicted the optimal parameters, providing a subject-specific description of the excitatory/inhibitory balance that could be analyzed at group level (as discussed above) or used to establish cluster membership in a data-driven approach.

While at group level there were noticeable differences in excitatory/inhibitory parameters, when considering single-subjects’ profiles, K-means clusters were not group specific; this suggests a heterogeneous excitatory and inhibitory balance across subjects that could be exploited for future personalized interventions. Considering the biophysical meaning of TVB parameters, cluster 1 was mainly characterized by overexcitation and was more frequent in healthy controls and Amyotrophic lateral sclerosis. As well as hyperexcitability is a well-known feature of Amyotrophic lateral sclerosis patients^12,67^, the presence of some healthy controls subjects in this cluster is not surprising. Indeed, the effect of aging on the glutamatergic system is currently under investigation^68^, and even if glutamate is mostly reported to decrease with age increase^69^, age-related effects on glutamatergic release and uptake processes and NMDA receptor activation could be differentially modulated in some healthy subjects. Even in cluster 2 we found some healthy controls, which presented not only high excitation but also high global coupling strength. This high G value could be due to an increased strength of long-range connectivity or increased synchrony between nodes. It is worth noting that cluster 2 was more common in pathological conditions than in healthy controls, and this is in line with the frequent observation of hyperexcitation and hypersynchrony in Alzheimer’s disease^11,65,70^, Frontotemporal Dementia^10,71^ and Amyotrophic lateral sclerosis^12,72^. Cluster 3 was the most related to healthy controls, while the number of patients was marginal (0 in Alzheimer’s disease). This cluster is mainly characterized by high recurrent excitation; interestingly, alterations of this property are less explored as potential mechanisms in clinical conditions. Only few network models have been developed to account for the influence of recurrent excitation and to explore its changes in pathologies. Strong self-excitation was shown to be required in network models to achieve satisfactory simulations of decision making and working memory tasks^73^, in line with our evidence of high recurrent excitation in healthy controls. Moreover, in a previously proposed computational model applied to Alzheimer’s disease the variation of local recurrent excitation has been suggested as a brain mechanism employed to compensate for alterations induced by other types of synapse loss^74^. Unfortunately, nothing is known about recurrent excitation in network models of Frontotemporal Dementia or Amyotrophic lateral sclerosis, but the presence of Frontotemporal Dementia and Amyotrophic lateral sclerosis patients in cluster 3 prompts to explore the effect of this parameter in clinical conditions other than Alzheimer’s disease. Finally, cluster 4 was mostly associated with Alzheimer’s disease patients, followed by Amyotrophic lateral sclerosis, Frontotemporal Dementia and healthy controls. While convergent findings are increasingly supporting the role of a GABA function increase in Alzheimer’s disease^13,75,76^, GABAergic dysfunction is mostly described as an overall decrease of cortical inhibition in Frontotemporal Dementia and Amyotrophic lateral sclerosis. Our results suggest the possibility of an increased GABAergic activity not only in Alzheimer’s disease patients, as already observed in literature, but also in subsets of subjects affected by other neurodegenerative conditions.

It is important to point out that Frontotemporal Dementia patients appeared to be the most distributed between clusters, without a main cluster membership, and this evidence reflects the heterogeneity of our Frontotemporal Dementia cohort, which is in line with the wide spectrum of neurotransmitters deficits which has been already observed in Frontotemporal Dementia^77^.

### Cerebellar role in neurodegeneration

#### Cerebellar impact in different clinical conditions

Cerebellar impairment has been consistently observed in neurodegenerative diseases, although for years the cerebellum has been rarely considered in neurodegenerative conditions. Disease-specific clusters of cerebellar atrophy have been found in Alzheimer’s disease, Frontotemporal Dementia and Amyotrophic lateral sclerosis^78^. Functional connectivity alterations^22,23^ and white matter disruption^79^ characterize cerebro-cerebellar loops in Alzheimer’s disease patients. Abnormal network connectivity between cerebellum and cerebral cortical regions has been confirmed in the main subtypes of Frontotemporal Dementia^80,81^ (behavioral-variant, semantic dementia and progressive nonfluent aphasia). In Amyotrophic lateral sclerosis the functional reorganization following motor neuronal loss increases cerebellar activation in motor tasks with respect to controls^26^, while a widespread pattern of white matter abnormalities has been reported together with white matter volume reduction^82^.

In our work, the integration of cerebro-cerebellar connections improved the predictive power of TVB simulations in pathological conditions, supporting cerebellar involvement in neurodegenerative states and confirming the sizeable contribution of cerebro-cerebellar connectivity to simulated brain dynamics ^16^. This improvement caused by the connection of cerebellar nodes in TVB was especially evident in Amyotrophic lateral sclerosis, which is a long-range motor neuron disease. This calls for future work to establish whether this result reflects a higher cerebellar recruitment determined by Amyotrophic lateral sclerosis functional reorganization^26^. Studies evaluating the impact of cerebellar integration not only on static FC simulations, as performed in this work, but also on dynamic resting-state FC simulations^83,84^ are warranted.

## Conclusion

TVB-derived biophysical parameters provided a unique description of the excitatory/inhibitory balance both at the group and single subject level. An extremely high performance was achieved in patients’ discrimination combining TVB parameters and neuropsychological scores. Moreover, the integration of cerebro-cerebellar connections in TVB improved the predictive power of the model in neurodegeneration. Overall, this work opens new perspectives for the use of TVB to explore neurodegenerative mechanisms, supports the involvement of the cerebellum in determining brain dynamics in neurodegenerative diseases, and suggests a novel approach to obtain physiological information relevant to future personalized diagnosis and therapy.

## Abbreviations

expFC: experimental Functional Connectivity
FC: Functional Connectivity
PCC: Pearson Correlation Coefficient
SC: Structural Connectivity
simFC: simulated Functional Connectivity
TVB: The Virtual Brain

## Supplementary material

**Supplementary Figure 1.**
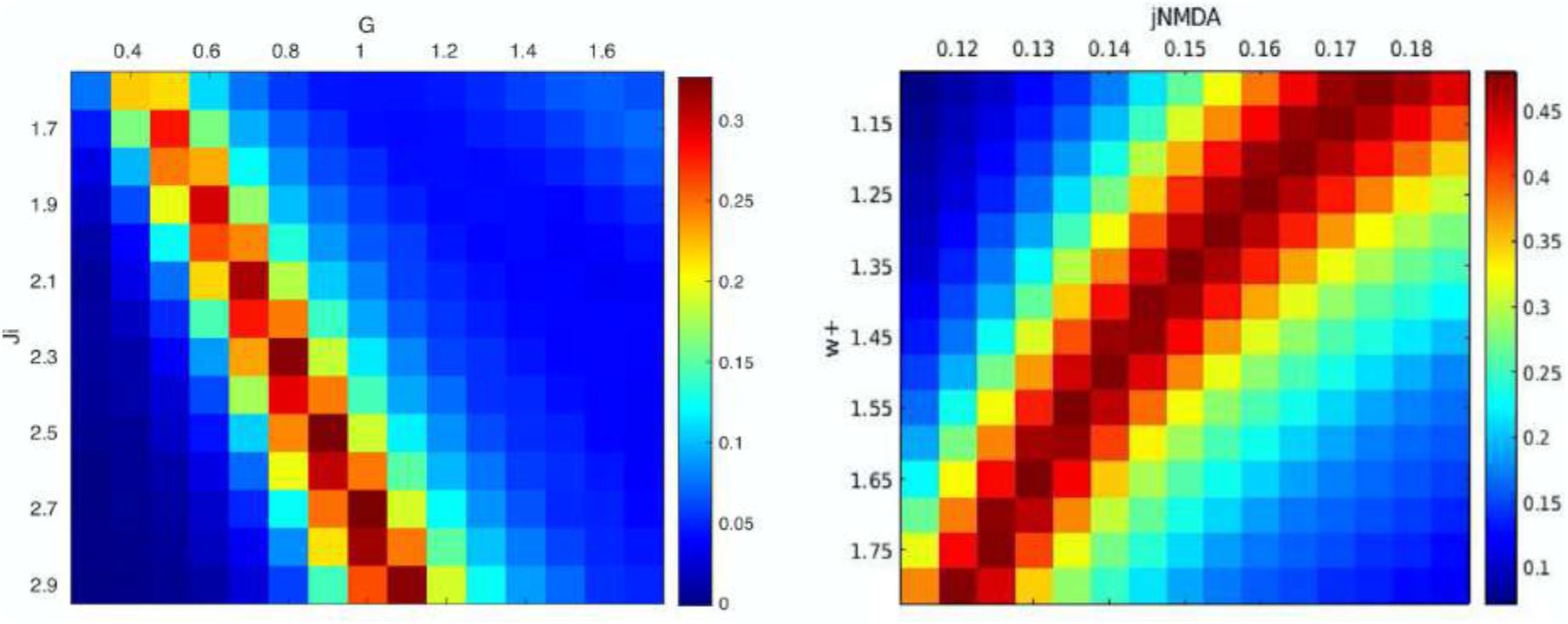
Parameter space exploration results. 2D parameter space heat maps show different values of correlation between experimental and simulated FC obtained at different combinations of the TVB parameters (G, Ji, J_NMDA and w+). Parameter values that yielded the highest correlation were chosen for the next steps of the analysis.

**Supplementary Figure 2.**
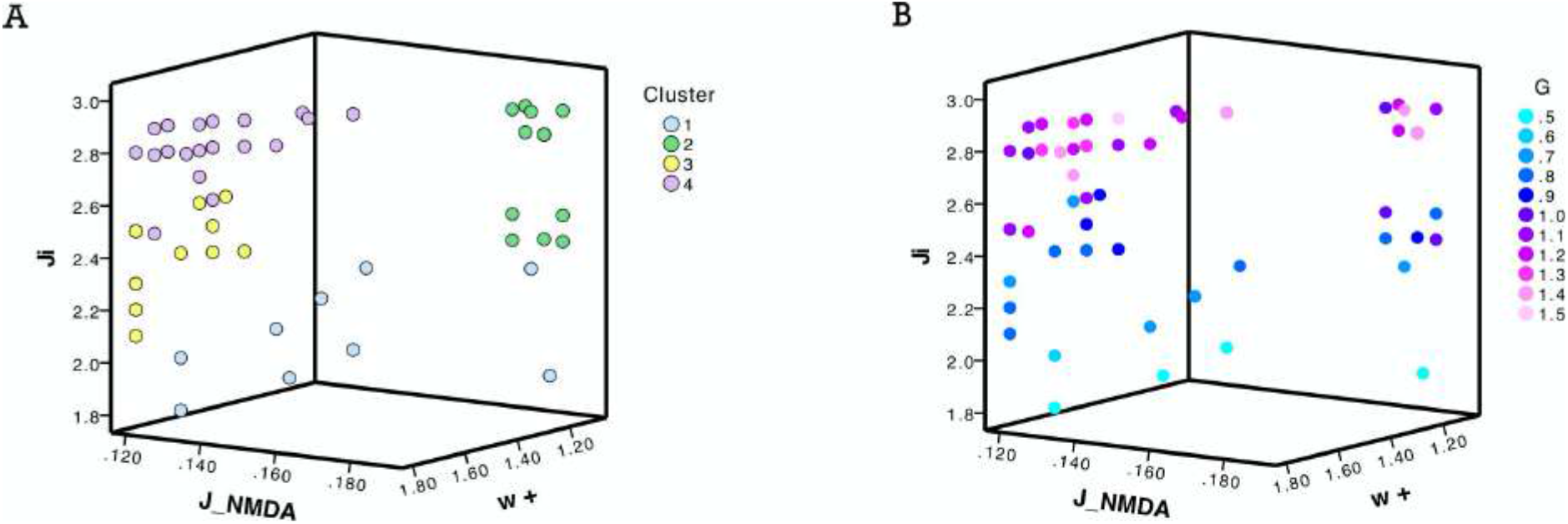
Clustering analysis. **(A)** Visual representation of the four clusters (in different colors) identified with k-means analysis using TVB-derived optimal biophysical parameters as input variables. **(B)** Each of the four clusters was characterized by a combination of low and high values of TVB-derived biophysical parameters (Ji, J_NMDA and wp are represented on the axis and G values are represented with color code).

**Supplementary Table 1:** Wong-Wang model parameters for TVB simulations.

| PARAMETERS | VALUE | DESCRIPTION |
| --- | --- | --- |
| $a_E, b_E, d_E, \tau_E, W_E$ | 310 nC<sup>-1</sup>, 125 Hz, 0.16 s, 100 ms, 1 | Excitatory gating variables |
| $a_I, b_I, d_I, \tau_I, W_I$ | 615 nC<sup>-1</sup>, 177 Hz, 0.087 s, 10 ms, 0.7 | Inhibitory gating variables |
| $\gamma$ | 0.641/1000 | Kinetic parameter |
| $\sigma$ | 0.01 nA | Noise amplitude |
| $I_0$ | 0.382 nA | Overall effective external input |
| $C_{ij}$ | Obtained from diffusion tractography | Structural connectivity (SC) matrix |
| $G$ | Obtained from parameters optimization | Global coupling scaling factor |
| $J_i$ | Obtained from parameters optimization | Feedback inhibitory synaptic coupling |
| $J_{NMDA}$ | Obtained from parameters optimization | Excitatory synaptic coupling |
| $w_+$ | Obtained from parameters optimization | Local excitatory recurrence |

**Supplementary Table 2:** Classification results for group comparisons.

|  | TVB_PARAMS |  |  | NPS |  |  | TVB + NPS |  |  |
| --- | --- | --- | --- | --- | --- | --- | --- | --- | --- |
|  | AUC | Sens. | Spec. | AUC | Sens. | Spec. | AUC | Sens. | Spec. |
| **HC vs AD** | 76.7% | 0.800 | 0.733 | 93.3% | 0.867 | 1.000 | 100% | 1.000 | 1.000 |
| **HC vs FTD** | 63.3% | 0.600 | 0.667 | 90% | 0.800 | 1.000 | 93.3% | 0.867 | 1.000 |
| **HC vs ALS** | 76.7% | 0.733 | 0.800 | 85.7% | 0.769 | 0.933 | 89.3% | 0.846 | 0.933 |
| **AD vs FTD** | 73.3% | 0.533 | 0.933 | 80% | 0.800 | 0.800 | 80% | 0.867 | 0.733 |
| **AD vs ALS** | 66.7% | 0.600 | 0.733 | 96.4% | 1.000 | 0.933 | 96.4% | 1.000 | 0.933 |
| **FTD vs ALS** | 56.7% | 0.533 | 0.600 | 96.4% | 1.000 | 0.933 | 100% | 1.000 | 1.000 |

Areas under the curve (AUC), sensitivity (Sens.) and specificity (Spec.) are reported for all classifications (HC=healthy controls, AD=Alzheimer's disease, FTD=Frontotemporal dementia, ALS=Amyotrophic lateral sclerosis) performed with different independent variables: TVB-derived biophysical parameters alone (TVB\_params), neuropsychological scores alone (NPS) and a combination of both (TVB+NPS).

